# Divergent Immediate and Delayed Effects of Juvenile Exposure to Doxorubicin on the Thymus in C57BL/6 Mice

**DOI:** 10.1101/2024.08.21.609003

**Authors:** B George, KJV Dahlquist, MKO Grant, MR Daniel, DM Smith, KT Sadak, D Seelig, CD Camell, BN Zordoky

## Abstract

**Background:** The understanding of alterations within the immune system following doxorubicin (DOX) chemotherapy, and subsequent restoration, in childhood cancer survivors remains limited. This investigation endeavors to elucidate the immediate and delayed changes in thymic immune cell populations and their phenotypes in response to clinically relevant low doses of DOX in a juvenile mouse model.

**Methods:** Male mice underwent a regimen of repeated low-dose DOX intraperitoneal injections at 4 mg/kg/week for three consecutive weeks. One week after the last dose of DOX, a subset of mice was euthanized to assess the immediate effects of DOX administration. A second subset of mice was euthanized five weeks after the last DOX dose to evaluate the delayed effects. Thymic samples were collected for multiparameter flow cytometry analysis to evaluate alterations in immune cell composition and phenotype. Additionally, quantitative real-time polymerase chain reaction (qRT-PCR) was employed to measure gene expression of^-^ cytokines and senescence markers.

**Results:** One week following DOX administration, DOX treatment resulted in significant decline in thymus weight, with notable alterations in immune cell subpopulations. Reduced frequencies of mature CD3^+^CD4^+^ and CD3^+^CD8^+^ T cells were observed, along with changes in proliferation and exhaustion markers. Gene expression analysis revealed upregulation of *Foxn*, *Pax1*, *Ifn*γ, and *Il7* alongside decreased *Il6* and *Il17* expression. Furthermore, *Cdkn1a* (*p21^Cip1^)* expression was elevated, suggesting immunosenescence. Five weeks following DOX administration, delayed effects of DOX treatment manifested in rebound increase in thymus weight and altered frequencies of CD4^+^ and CD8^+^ T cell subsets, with distinct patterns of proliferation and exhaustion observed. Notably, central memory CD4^+^ T cells exhibited significant decrease in frequency, while naïve and effector memory CD4^+^ T cells showed reduced proliferation (Ki67^+^) and PD1 expression. Similar trends were observed in CD8^+^ T cell subsets, indicating selective effects of DOX on T cell differentiation and function. Although expression of thymus-related genes was normalized, *p21^Cip1^* gene expression remained elevated.

**Conclusion:** DOX treatment elicits a multifaceted influence on immune cell subsets and thymic weight. Immediate effects included thymic atrophy and reductions in mature T cell populations, while delayed effects showed rebound thymic hyperplasia and selective changes in CD4^+^ and CD8^+^ T cell subsets. Notably, both central memory and effector memory T cells exhibited reduced proliferation and exhaustion, suggesting unique impacts of DOX on immune cell function. The enduring elevation in *p21^Cip1^* gene expression 5 weeks after DOX treatment suggests an immunosenescent phenotype. These observations collectively illuminate the formidable task of preserving immune competence and overall well-being in childhood cancer survivors subjected to DOX therapy.

## Introduction

Doxorubicin (DOX) chemotherapy, a widely employed treatment for cancer, has been shown to cause a transient suppression of immune cell function and heightened vulnerability to infections among cancer patients [1]. The thymus plays a vital role as the site where T lymphocytes (T cells), a subset of immune cells central in the response against infectious pathogens and cancer, develop and mature. While the thymus is most active during childhood and adolescence, it diminishes in size and activity with age, exhibiting signs of immunosenescence [2]. Chemotherapy can also cause deleterious effects on this organ [1, 3]. The administration of chemotherapy induces thymic atrophy, leading to a reduction in the generation of naïve T cells and compromised cellular immunity function [4–6]. Particularly, DOX chemotherapy impacts rapidly dividing cells, thus thymocytes, which undergo a massive proliferative burst within the thymus, would be highly affected. The suppression of thymic function leads to a decline in the production and maturation of T cells, thereby influencing immune cell homeostasis and the immune response to infectious challenge [7, 8]. When the thymus undergoes atrophy, the production of mature naive T cells is compromised [9]. Research has shown that DOX, a chemotherapy drug used to treat various cancers, can induce thymic atrophy [10]. This means that exposure to DOX can lead to the shrinking or reduction in size of the thymus gland, potentially affecting immune function by disrupting T-cell production. However, upon discontinuation of chemotherapy, thymic atrophy may undergo recovery, potentially giving rise to a hyperplastic process known as rebound thymic hyperplasia [11–13]. Thymic hyperplasia is associated with robust thymic regeneration, characterized by an increase in both thymic size and density, concomitant with the restoration of thymic T cell output [4,13,14]. Although this phenomenon is common in children and adolescents and occasionally observed in young adults, it is rare in older patients [15–18].

T cells are immune cells that play a central role in the immune system. T cells arise from hematopoietic stem cells differentiating into lymphoid progenitors within the bone marrow [19]. Within the thymus, these lymphoid progenitors are called thymocytes and undergo a maturation process to generate mature T cells with rearranged, highly specific, T cell receptors (TCR). Initially, thymocytes are double negative for CD4 and CD8 and do not express the TCR. They then begin to express TCR and become double positive for CD4 and CD8 where they then undergo positive and negative selection. In the last maturation step within the thymus, mature naïve T cells are generated as they become single positive for either CD4 or CD8. Once mature, naïve T cells then circulate in the blood and lymph enter circulation to encounter their specific antigen [20]. CD8^+^ T cells are cytotoxic and aid in the immune response by secreting effector cytokines such as tumor necrosis factor alpha (TNF-α) and interferon-gamma (IFN-γ) and directly killing pathogenic cells through cytotoxic mechanisms [21]. CD4^+^ T cells can be divided further into either Tregs or T helper cells that have specialized roles in supporting a specific immune response [21]. Once T cells encounter their specific antigen they then differentiate into memory T cells [22]. Central memory T cells are long lasting, remaining for years after initial antigen encounter, whereas effector memory T cells are potent responders and are short lived [11]. Alterations in the development and phenotype in these cells can lead to alterations in the immune response to infections and cancer. Both atrophy and hyperplasia of the thymus can impact T cell development and function, potentially affecting immune responses to cancer and other diseases [23].

The enduring impacts of cancer treatment extend to prolonged immune suppression and cellular senescence, persisting for over a decade post-treatment [27, 28]. Studies reveal that cancer survivors continue to undergo lasting alterations in their immune profiles well after chemotherapy completion [29-31]. These changes involve reduced T cell counts, impaired T cell function, and disruptions in immune checkpoint regulation. Additionally, chemotherapy-induced cellular senescence, characterized by permanent cell cycle arrest and functional changes, significantly influences aging and age-related diseases, including cancer [29]. The induction of senescence in both cancerous and healthy cells, including immune cells, underscores its role in aged-related outcomes. Such immune system modifications heighten susceptibility to infections and weaken the body’s ability to detect and combat cancer recurrence. The present study investigates the intricate and multifaceted impact of DOX chemotherapy on immune cell subpopulations and thymic functionality in juvenile male mice, focusing on both immediate and delayed repercussions. While previous studies have largely focused on the immediate cytotoxic impacts of chemotherapy [8], there remains a significant gap in understanding its specific influence on the immune system, especially under the conditions of repeated low-dose treatments. This research extends into the relatively unexplored domain of chemotherapy-induced immune modulation by conducting a thorough analysis of both the immediate and longer-term immune responses to DOX.

## Materials and Methods

### Animals

All animal-related experimental procedures were ethically approved by the University of Minnesota Institutional Animal Care and Use Committee (IACUC protocol number: 2106-39176A). The mice were housed in a controlled environment with specific pathogen-free (SPF) conditions, maintained in a 14-hour light/10-hour dark cycle at a temperature of 21 ± 2°C, and provided with ad libitum access to food and water. To ensure acclimatization, mice were introduced to the animal facility one week prior to the commencement of experimental procedures. The overall experimental designs are schematically represented in Fig. 1A and Fig. 5A. Commencing at five weeks of age, mice received intraperitoneal injections of DOX at a dosage of 4 mg/kg/week for three consecutive weeks, or an equivalent volume of sterile saline, resulting in a cumulative dose of 12 mg/kg. Weekly assessments of animal weights were conducted, and subsets of mice were humanely euthanized one week after the final dose and another subset after a five-week interval. Euthanasia was performed in a humane manner using decapitation under isoflurane anesthesia. Subsequently, thymic tissues were harvested, cleansed under a microscope in ice-cold phosphate-buffered saline solution, and either promptly flash-frozen in liquid nitrogen or placed in cold media for immune cell isolation and analysis. The frozen specimens were stored at -80°C for subsequent analytical procedures.

**Figure 1:**
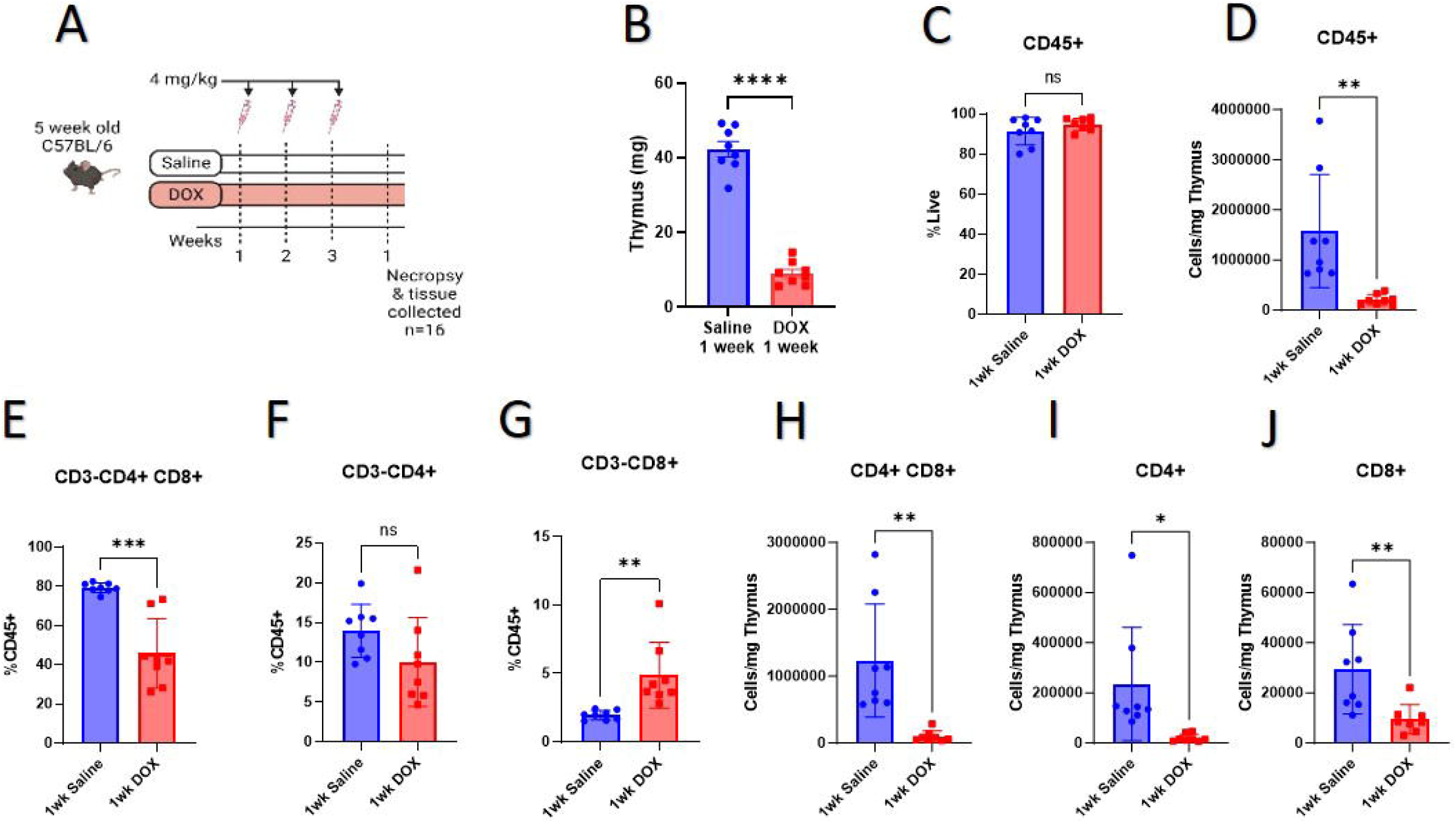
DOX induced changes in thymus weight and T cell populations one week after treatment: **A)** Experimental design to assess the immediate effects of DOX-induced immune dysfunction. Male mice were exposed to a series of low-dose DOX administrations at 4 mg/kg/week for three consecutive weeks. Mice were euthanized one week after the final treatment to observe immediate effects of DOX. Thymus samples were collected for flow cytometry analysis; **B)** Immediate effects of DOX on thymus weights (mg); Bar graphs representing the **C)** frequency and **D)** cell count (per milligram (mg) of thymus) of immune cells (CD45^+^); Bar graphs representing the frequencies of **E)** CD4^+^CD8^+^, **F)** CD4^+^, **G)** CD8^+^ and cell count of **H)** CD4^+^CD8^+^, **I)** CD4^+^ and **J)** CD8^+^ cells per mg of thymus;. n = 8 mice/group. \**p* < 0.05, \*\**p* < 0.01, \*\*\**p* < 0.001, and \*\*\*\**p* < 0.0001 vs. saline control group (analyzed by unpaired t-test).

### Flow and Antibodies

#### Immune cell isolation and staining

Immune cells were isolated from the thymus using mechanical digestion. Red blood cells were lysed using ACK lysis buffer. For staining, cells were stained with a fixable viability dye for 25 min at 4°C in the dark. Cells were then incubated in Fc block (anti-CD16 & anti-CD32) and surface antibodies for 45 min at 4°C protected from light. For intracellular or nuclear staining, cells were fixed and permeabilized using the BD Cytofix/Cytoperm kit (554715) or eBioscience FOXP3/Transcription factor staining buffer set followed by nuclear antibodies staining for 45 min at 4°C protected from light. All antibodies used were confirmed to have been validated by their manufacturer. Flow cytometry data were acquired on a BD FACSymphony A3 Cell Analyzer R6609 and analyzed using Flowjo software version 10. Gating strategies are provided in Sup. Fig. 2 and 3.

#### RNA Extraction and Quantitative Real-Time PCR

Total RNA was isolated from cryopreserved thymus tissues utilizing Invitrogen TRIzol® reagent (Thermo Fisher Scientific, Waltham, MA, USA), adhering to the manufacturer’s guidelines. The quantification of RNA concentrations was executed at 260 nm employing a NanoDrop 8000 spectrophotometer (Thermo Fisher Scientific). Subsequently, first-strand cDNA was synthesized from 1.5 μg of total RNA, employing the Applied Biosystems high-capacity cDNA reverse transcription kit (Thermo Fisher Scientific), following the manufacturer’s instructions. Quantitative real-time polymerase chain reaction (qRT-PCR) was employed for the quantification of specific mRNA expression, involving PCR amplification of the synthesized cDNA within 384-well optical reaction plates using an Applied Biosystems QuantStudio 5 instrument (Thermo Fisher Scientific). The 20 µL reaction mixture consisted of 1 µL of the cDNA sample, 0.025 µL of 30 µM forward primer, and 0.025 µL of 30 µM reverse primer (attaining a final concentration of 40 nM for each primer). Additionally, it included 10 µL of Applied Biosystems SYBR Green Universal Mastermix (Thermo Fisher Scientific) and 8.95 µL of nuclease-free water. The thermocycler conditions comprised an initial denaturation step at 95 °C for 10 minutes, succeeded by 40 PCR cycles involving denaturation at 95 °C for 15 sec and annealing/extension at 60 °C for 1 minutes. Gene expression analyses were conducted using primers sourced from previously published studies, detailed in Table 1. Normalization of mRNA expression levels was performed relative to *Actb*, and the relative expressions were determined utilizing the ΔΔCt method.

**Table 1.**
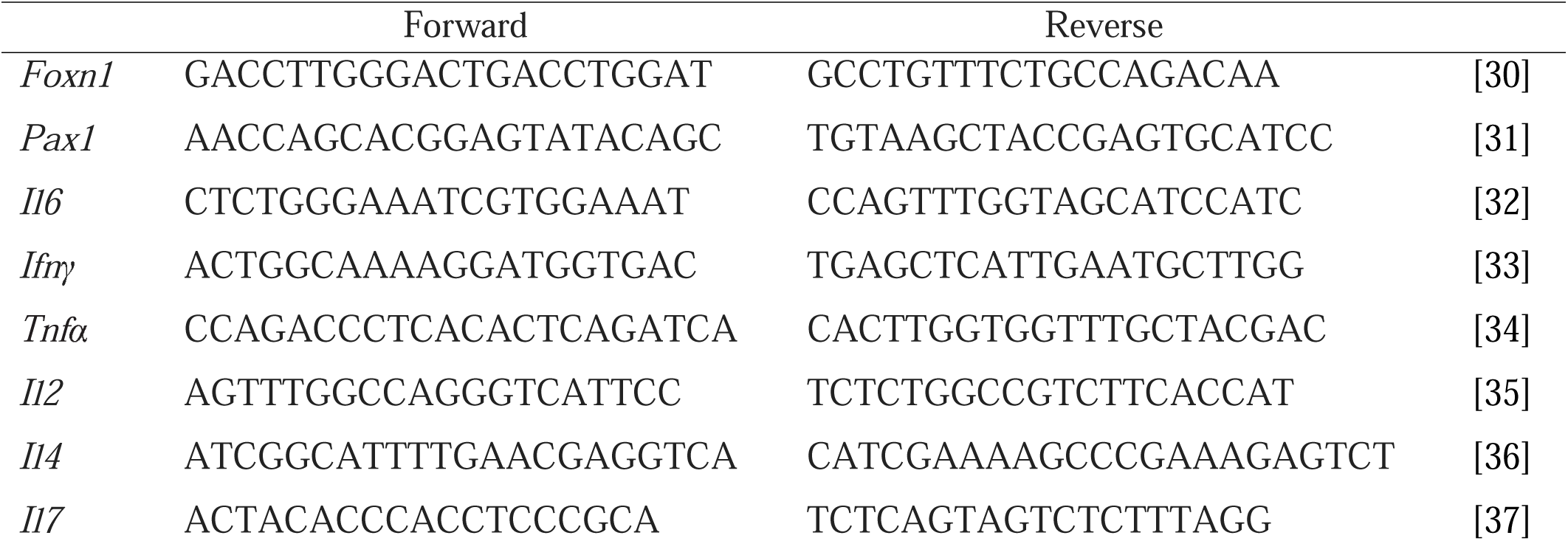
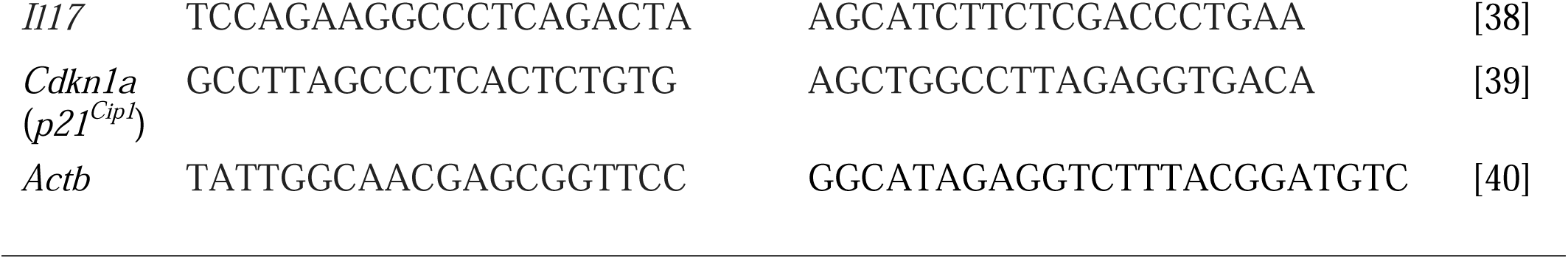
Primer sequences used for real-time PCR experiments.

#### Histopathological Analysis

The tissues were fixed in 10% neutral buffered formalin, transferred to 70% ethanol, and processed into paraffin blocks. From each paraffin block, a single 4 µm section was cut and stained with hematoxylin and eosin. All samples were evaluated by a single, board-certified veterinary pathologist (DMS) that was blinded to study group information. Each sample was evaluated for hemorrhage, necrosis, inflammation, edema, and cystic degeneration with each finding semi-quantitatively scored 0-4 according to the following rubric: 0 = finding identified, 1 = finding is rare and represents <5% of the total tissue surface area, 2 = mild severity and affects 5-25% of the tissue surface area, 3 = moderate severity and affects 26-50% of the tissue surface area, and 4 = marked severity and affects >51% of the tissue surface area.

#### Statistical Analysis

Data is presented as mean ± standard error of the mean (SEM). To assess differences between treatment groups, unpaired Student’s two-tailed t-test was conducted. Statistical significance was established at a probability value of < 0.05. Analysis was done in GraphPad Prism 9.5.1.

## Results

The current study aimed to determine the direct effects of DOX on the thymus in young male mice, specifically examining changes in immune cell numbers and proportions. Male mice were subjected to a regimen of repeated low-dose DOX administration at 4 mg/kg/week for three consecutive weeks (Fig 1A). To maintain clinical relevance, we delivered an intentionally low level of DOX, 4 mg/kg/week, equivalent to approximately ∼40 mg/m^2^ in humans [41]. Our prior work has established the absence of severe morbidity or any mortality when administering 4 mg/kg/week for 3-5 weeks [44, 45]. One week after the last administration of DOX, mice were euthanized to evaluate the immediate effects of DOX exposure. Notably, no mortality incidents were recorded following the third DOX dose. Male mice subjected to DOX showed a significant decline in body weight 1 week after the final injection compared to saline treated mice (Sup. Fig. 1A and 1B).

In DOX-treated male mice, we investigated the immediate impact of DOX on thymus weight, which revealed a significant reduction in weight compared to the saline group (Fig. 1B). The decrease in weight was maintained when normalized to tibial length and or to body weight, indicating that organ specific effects were occurring (Sup. Fig. 1C and 1D). To determine the potential cellular source of thymic atrophy 1 week after DOX treatment we identified total immune cell (CD45^+^) frequencies and cells per mg of tissue. There were no differences in the frequency of total immune cells, but there was a marked reduction in the overall number of immune cells per mg of thymus (Fig. 1C and 1D) in the DOX group when compared to the saline group. We used multi-parameter flow cytometry to examine developing thymocytes (Sup. Fig. 2 and 3). We observed a reduced frequency of double positive (CD4^+^CD8^+^) cells in the DOX mice compare to saline control one week after the last DOX treatment (Fig. 1E). No significant difference was observed in the single positive, CD4^+^ cell frequency but an increase was seen in the single positive, CD8^+^ cell frequency in the DOX group when compared to the saline group (Fig. 1F and 1G). Furthermore, the cells per mg of thymus of double positive (CD4^+^ CD8^+^), CD4^+^, and CD8^+^ cells, were significantly reduced in DOX treated mice compared to those given saline (Fig. 1H, 1I, and 1J). These results are consistent with previous work that describes the atrophy of the thymus following DOX-treatment due to reduced thymocyte cell numbers [46, 47].

The marker Ki67 is used to define proliferating cells as it is expressed only by proliferating cells [46]. To address the phenotype of mature CD3^+^ T cells, we analyzed naïve and memory status, as well as markers of proliferation (Ki67) and exhaustion (PD1) [47] ( Sup. Fig. 5A). The frequency of mature CD3^+^CD4^+^ T cells did not exhibit statistical significance between the saline and DOX-treated groups (Fig. 2A). There were no differences in the frequency of naïve CD4^+^ T cells (CD44^-^CD62^+^) and their proliferation, as indicated by Ki67 expression, between the two groups (Fig. 2B and 2E). However, there were significantly elevated frequencies of exhaustion, as assessed by PD1 expression, in the naïve CD4 single positives within the DOX-treated group (Fig. 2G). Upon encountering antigen, naïve T cells can acquire a central memory (CD44^+^CD62^+^) or effector memory (CD44^+^CD62^-^) phenotype. The frequencies of central and effector memory populations were unchanged when comparing saline to DOX mice (Fig. 2C and 2D). Ki67^+^ and PD1^+^ effector memory CD4^+^ T cells were increased in the DOX mice compared to the saline group (Fig. 2F and 2H).

**Figure 2:**
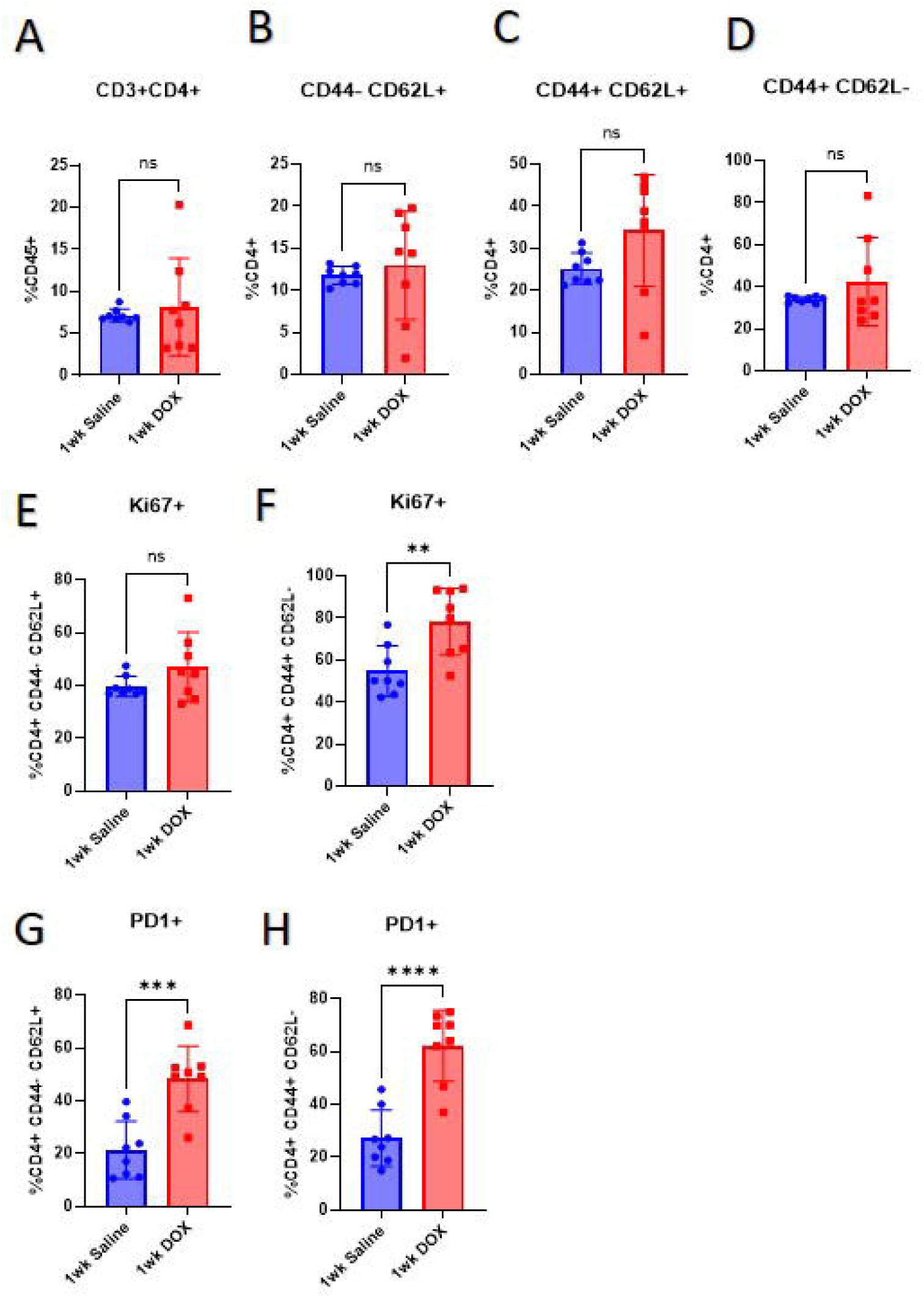
DOX induced changes in CD4^+^ T cell populations one week after treatment: Bar graphs representing frequencies of **A)** mature CD3^+^CD4^+^, **B)** naïve CD4^+^ T cells (CD44^-^ CD62L^+^), **C)** central memory CD4^+^ T cells (CD44^+^CD62L^+^) and **D)** effector memory CD4^+^ T cells (CD44^+^CD62L^-^);Bar graphs representing frequencies of **E)** Ki67^+^ in naïve CD4^+^ T cells and **F)** Ki67^+^ in effector memory CD4^+^ T cells; Bar graphs representing frequencies of **G)** PD1^+^ in naïve CD4^+^ T cells and **H)** PD1^+^ in effector memory CD4^+^ T cells. n = 8 mice/group. ns *p* > 0.05, \**p* < 0.05, \*\**p* < 0.01, \*\*\**p* < 0.001 and \*\*\*\**p* < 0.0001vs. the saline control group (analyzed by unpaired t-test).

We next examined the mature CD3^+^CD8^+^ T cells in the saline and DOX-treated groups. There was no significant difference in the frequency of mature CD3^+^CD8^+^ T cells within the DOX group but there was a reduced frequency of naïve CD8^+^ T cells (Fig. 3A and 3B). Although the frequency of proliferative naïve CD8^+^ T cells (Fig. 3E) showed no significant change, there was an elevated frequency of PD1^+^ naïve CD8^+^ T cells within the DOX group (Fig. 3H). DOX induced significantly increased frequencies of central memory CD8^+^ T cells but no significant differences in effector memory CD8^+^ T cells (Fig. 3C and 3D). The frequencies of Ki67^+^ central memory and effector memory CD8^+^ T cells increased in the DOX group compared to saline control (Fig. 3F and 3G). Additionally, PD1^+^ central and effector memory T cells increased with DOX treatment indicating an exhausted phenotype (Fig. 3I and 3J). These results are consistent with DOX-induced alterations of T cell subsets including an increase in exhausted T cells.

**Figure 3:**
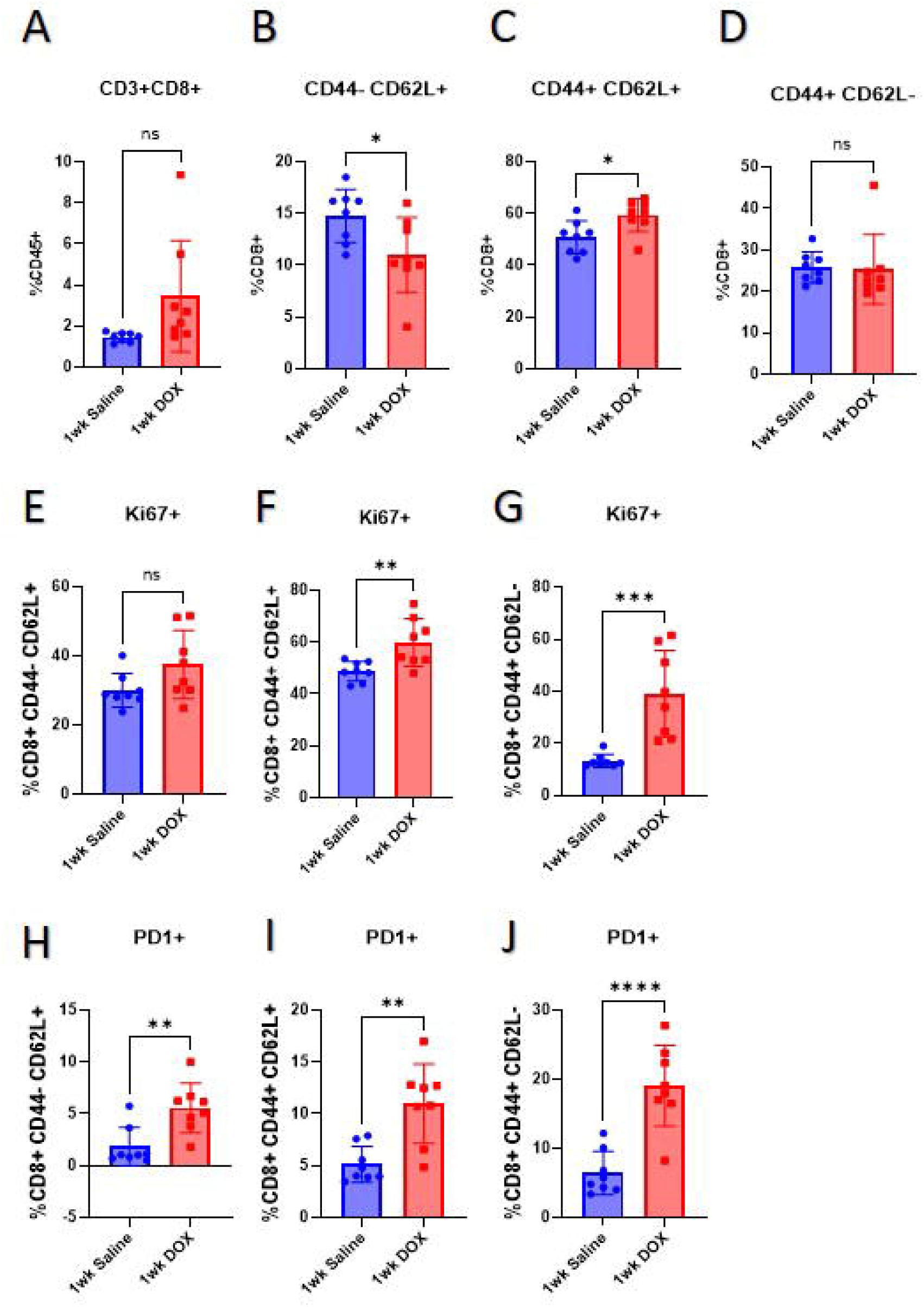
DOX induced changes in CD8^+^ T cell populations one week after treatment: Bar graphs representing frequencies of **A)** mature CD3^+^CD8^+^, **B)** naïve CD8^+^ T cells, **C)** central memory CD8^+^ T cells and **D)** effector memory CD8^+^ T cells; Bar graphs representing frequencies of **E)** Ki67^+^ in CD8^+^ naïve T cells, **F)** Ki67^+^ in central memory CD8^+^ T cells and **G)** Ki67^+^ in effector memory CD8^+^ T cells; Bar graphs representing frequencies of **H)** PD1^+^ in naïve CD8^+^ T cells, **I)** PD1^+^ in central memory CD8^+^ T cells and **J)** PD1^+^ in effector memory CD8^+^ T cells. n = 8 mice/group. ns *p* > 0.05, \**p* < 0.05, \*\**p* < 0.01, \*\*\**p* < 0.001 and \*\*\*\**p* < 0.0001 vs. saline control group (analyzed by unpaired t-test).

### Immediate Impact of DOX Administration on Thymic Gene Expression

Foxn1 serves as a central orchestrator of the intricate developmental processes governing the lineage of thymic epithelial cells, facilitating the downstream transcriptional activation of genes essential for thymus organogenesis and the comprehensive differentiation of thymic epithelial cells [48]. Additionally, PAX1 expression within the thymic epithelium is indispensable for establishing a specialized microenvironment crucial for normal T cell maturation [49]. In mice treated with DOX, an increase in *Foxn1* and *Pax1* expression was observed, indicating potential alterations in thymic development and T cell maturation (Fig. 4A and 4B). The cytokine network within the thymus involves dynamic interactions between various cytokines such as interleukin-6 (IL-6), interferon-gamma (IFN-γ), transforming growth factor-beta, IL-1, and IL-8 [50–52]. The immediate impact after DOX treatment in male mice resulted in decreased gene expression of *Il6* (Fig. 4C). DOX treatment is associated with thymic degeneration and senescence, leading to thymic atrophy and potential immunosuppression [10, 53]. We observed an increase in *p21^Cip1^* gene expression (Fig. 4G) and no differences in other senescence markers (Sup. Fig. 5) in the thymus one week following DOX administration.

**Figure 4:**
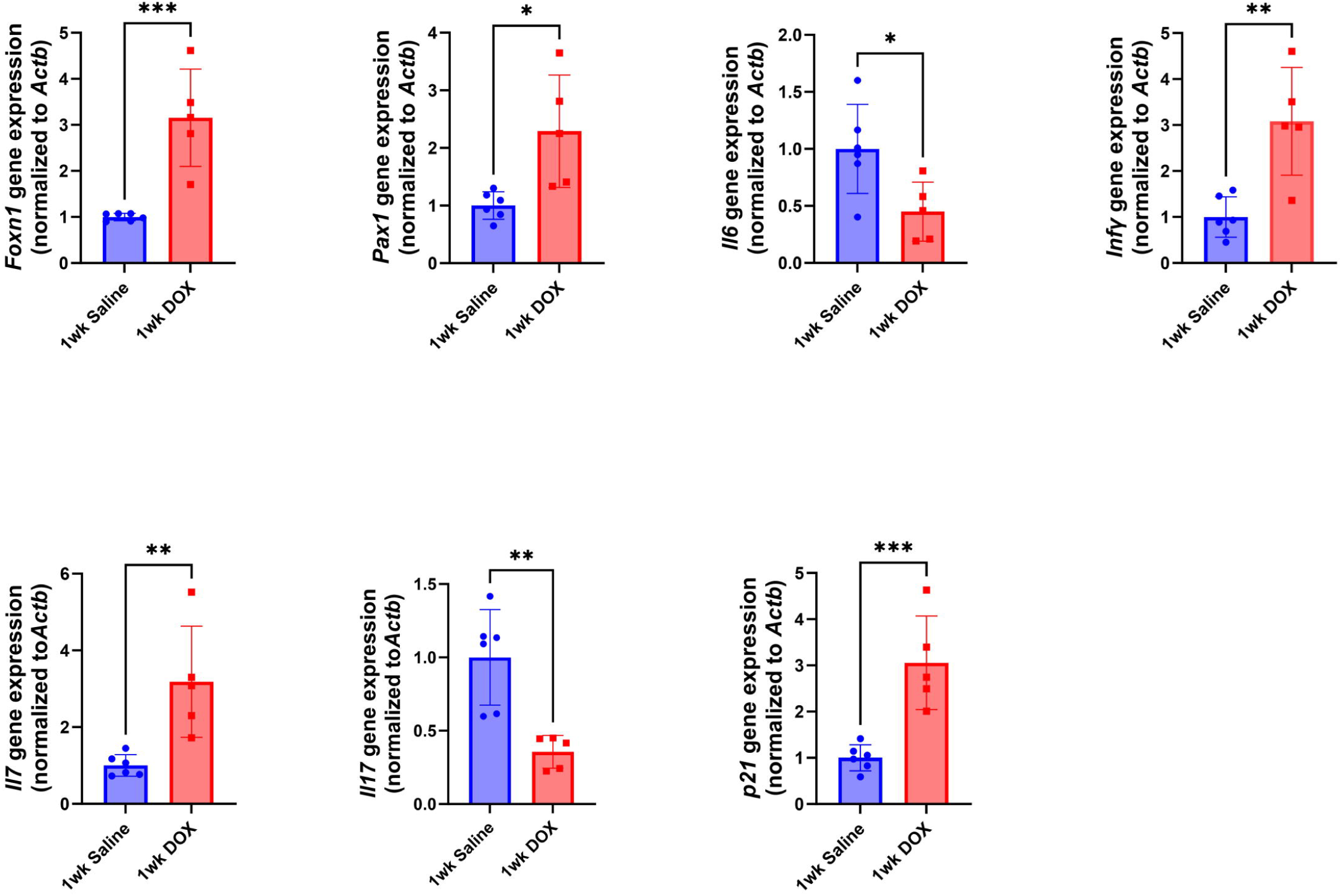
Immediate effects of DOX on the cytokines and senescence marker in thymus: Thymus were harvested from male mice one weeks following the administration of 4 mg/kg/week DOX or saline for three weeks (n = 5–8 per group). Following the extraction of total RNA, the mRNA expression of **A)** *Foxn1,* **B)** *Pax1,* **C)** *Il6*, **D)** *Ifn*γ **E)** *Il7*, **F)** *Il17*, and **G)** *p21^Cip1^* was determined by qRT-PCR. Values were normalized to ACTB and expressed relative to saline-treated male mice. Values are shown as the means ± SEMs. The statistical significance of pairwise comparisons was determined by unpairedt-test (\**p* < 0.05, \*\**p* < 0.01, and \*\*\**p* < 0.001.Furthermore, IFN-γ, an immunoregulatory cytokine, modulates the proliferation and differentiation of immunologically active cells, promoting the expansion of CD4^+^ T-helper type 1 (Th1) cells while inhibiting Th2 cell growth [53]. Interestingly, DOX treatment led to an increase in *Ifn*γ expression (Fig. 4D). We also observed an increase in *Il7,* a critical cytokine for T cell survival, in the DOX-treated mice (Fig. 4E). In contrast, there was a decrease in *Il17* expression in DOX-treated mice (Fig. 4F). Therefore, in the context of DOX treatment, a decrease in *Il17* expression may contribute to thymic atrophy, which can impact T cell development and immune regulation. DOX did not alter *Il2, Tnf*α and *Il4* one week following DOX administration (Sup. Fig. 4).

### Delayed effects of DOX on the thymus

Juvenile male mice underwent a regimen of repeated low dose DOX administrations at 4 mg/kg/week for three consecutive weeks. Subsequently, a five-week period without drug administration was introduced to assess the delayed effects of DOX exposure (Fig 5A). Final body-weigh was not different between saline and DOX-treated mice (Sup. Fig. 6A and 6B). In contrast to the immediate impact of DOX-induced reduction in body weights (Fig. 1B and Sup. Fig. 1A), the delayed impact of DOX on thymus weight revealed a notable increase in weight compared to the saline group (Fig. 5B). This increase in thymus weight was maintained when normalized to tibial length or to body weight (Sup. Fig. 6C and 6D).

**Figure 5:**
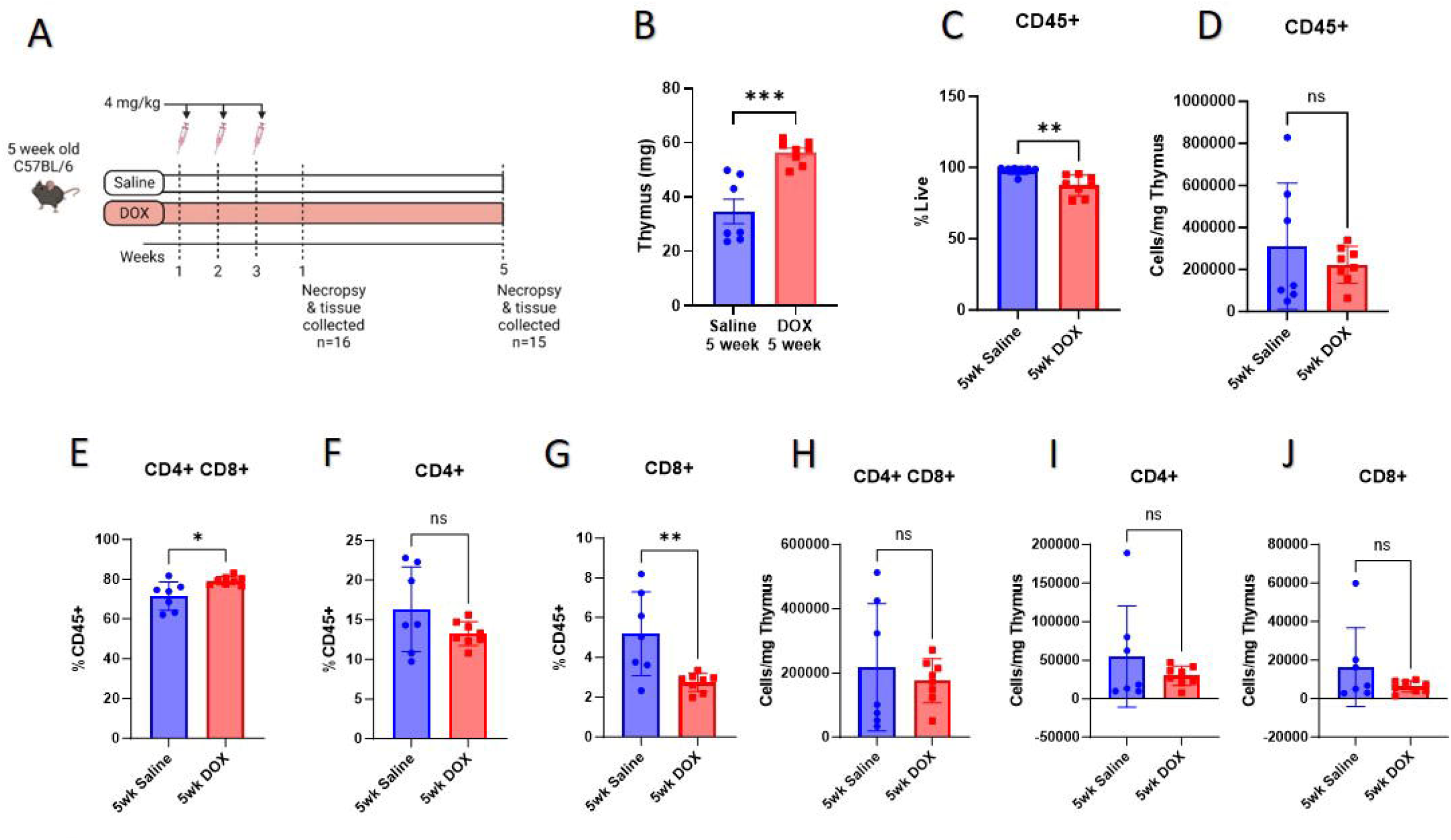
DOX induced changes in thymus weight and T cell populations five week after treatment: **A)** Experimental design to assess the delayed effects of DOX-induced immune dysfunction. Male mice were exposed to a series of low-dose DOX administrations at 4 mg/kg/week for three consecutive weeks. Eight DOX-treated and seven saline-treated mice were euthanized five weeks after the final treatment to observe the delayed effects of DOX. Thymus samples were collected for flow cytometry analysis; **B)** delayed effects of DOX on the thymus weight (mg); Bar graphs representing the **C)** frequency and **D)** cell count of hematopoietic cells (CD45^+^); Bar graphs representing the frequencies of **E)** double-positive i.e., CD4^+^CD8^+^, **F)** CD4^+^ T cells, **G)** CD8^+^ T cells and cell count of **H)** double^-^positive CD4^+^CD8^+^, **I)** CD4^+^ and **J)** CD8^+^ cells (in per mg thymus), five-week DOX group compared with the five-week saline group. n = 8 mice/group. ns *p* > 0.05 vs. the one-week saline control group (analyzed by unpaired t-test).

To address the possibility that the immediate effects of DOX are maintained to the five^-^week timepoint, we quantified the frequency and numbers of developing thymocytes. There was a diminished frequency of immune cells (CD45^+^), with no difference in cells/mg thymus at 5 weeks after DOX administration (Fig. 5C and 5D). There was a higher frequency of double-positive CD4^+^CD8^+^ cells (Fig. 5E). Although, there were no alterations in the frequency of CD4^+^ cells, there was a notable decrease in the frequency CD8^+^ T cells in the DOX group compared to the saline group (Fig. 5F and 5G). No differences were found in the double-positive CD4^+^CD8^+^, CD4^+^ cells, and CD8^+^ cells in the cells/mg of thymus (Fig. 5H, 5I, and 5J).

We first examined the mature CD4 T cell subset in the thymus. We detected a reduction in the frequency of mature CD3^+^CD4^+^ cells at five weeks following DOX treatment (Fig. 6A); however, there were no differences in the frequency of naïve CD4^+^ T cells (CD44^-^CD62L^+^) (Fig. 6B), a small, but significant decrease in the central memory CD4^+^ T cells (CD44^+^CD62L^+^) (Fig. 6C) and no difference in effector memory CD4^+^ T cells (CD44^+^CD62L^-^) (Fig. 6D). Ki67^+^ and PD1^+^ naïve CD4^+^ T cells were significantly decreased in the DOX-treated group compared to the saline group (Fig. 6E and 6G) These findings suggest that DOX may exert a suppressive effect on the activation and functional exhaustion of naïve CD4^+^ T cells. The frequencies of Ki67^+^ and PD1^+^ central memory CD4^+^ T cells were significantly lower in the DOX-treated group compared to the saline group (Fig. 6F and 6H).

**Figure 6:**
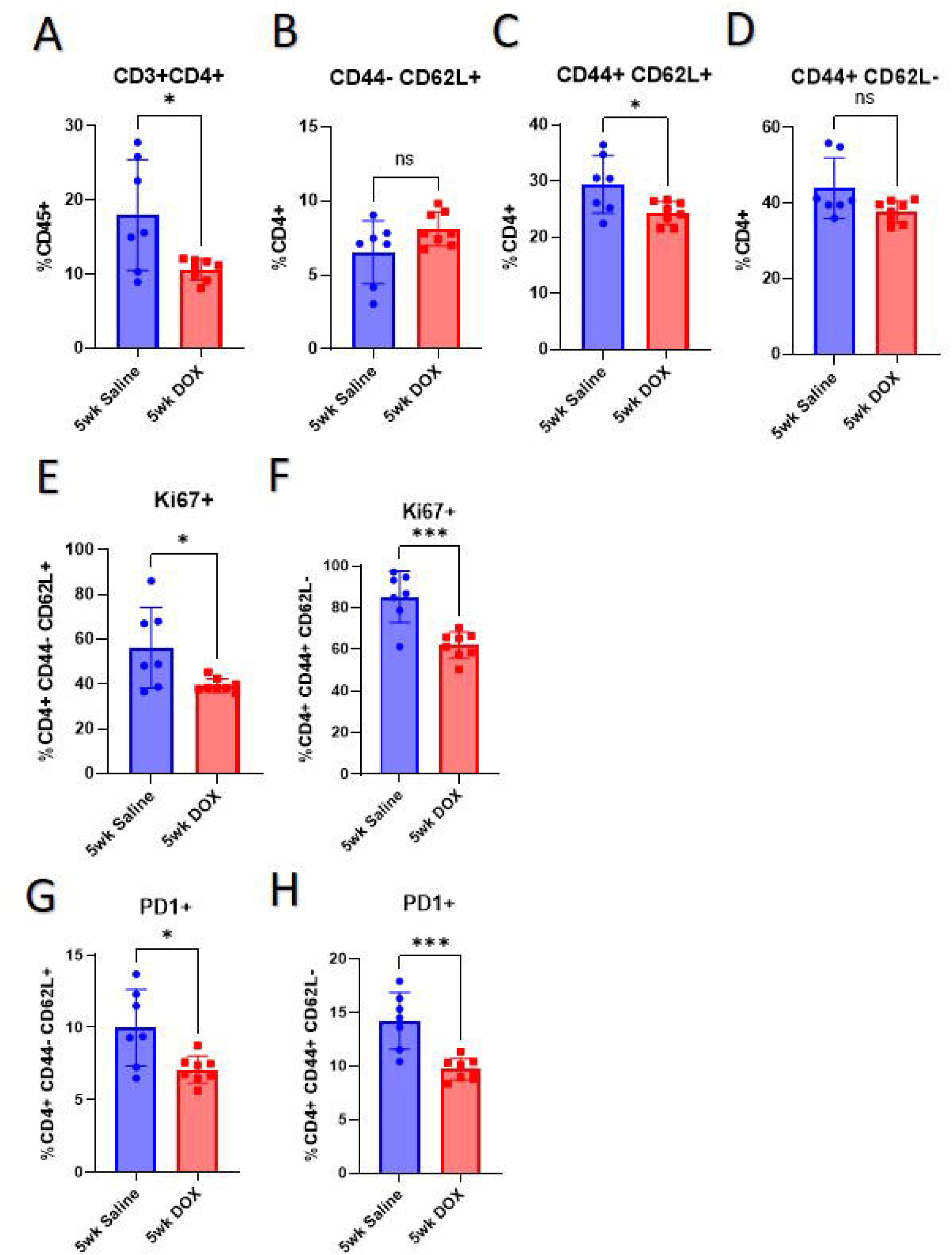
DOX induced changes in CD4^+^ T cell populations five week after treatment: Bar graphs representing frequencies of **A)** mature CD3^+^CD4^+^, **B)** naïve CD4^+^ T cells (CD44^-^ CD62L^+^), **C)** central memory CD4^+^ T cells (CD44^+^CD62L^+^) and **D)** effector memory CD4^+^ T cells (CD44^+^CD62L^-^); Bar graphs representing frequencies of **E)** Ki67^+^ in naïve CD4^+^ T cells and **F)** Ki67^+^ in effector memory CD4^+^ T cells; Bar graphs representing frequencies of **G)** PD1^+^ in naïve CD4^+^ T cells and **H)** effector memory CD4^+^ T cells; n = 7-8 mice/group. ns *p* > 0.05, \**p* < 0.05 and \*\*\**p* < 0.001 vs. the saline control group (analyzed by unpaired t-test).

We next examined the mature CD8 T cell subsets in the thymus from saline and DOX-treated mice. The mature CD3^+^CD8^+^ T cell population displayed a significant decrease (Fig. 7A). Conversely, the subsets of naïve CD8^+^ T cells exhibited a notable increase (Fig. 7B). The central memory and effector CD8^+^ T cells showed no significant changes in their frequencies following DOX treatment (Fig. 7C and 7D). There was a reduction in the frequency of proliferating naïve, central memory, and effector memory CD8^+^ T cell subsets in the DOX group compared to the saline group (Fig. 7E, 7F, and 7G). Additionally, there were decreases in PD1^+^ frequencies in the DOX group in naïve, central memory, and effector memory CD8^+^ T cells (Fig. 7H, 7I, and 7J).

**Figure 7:**
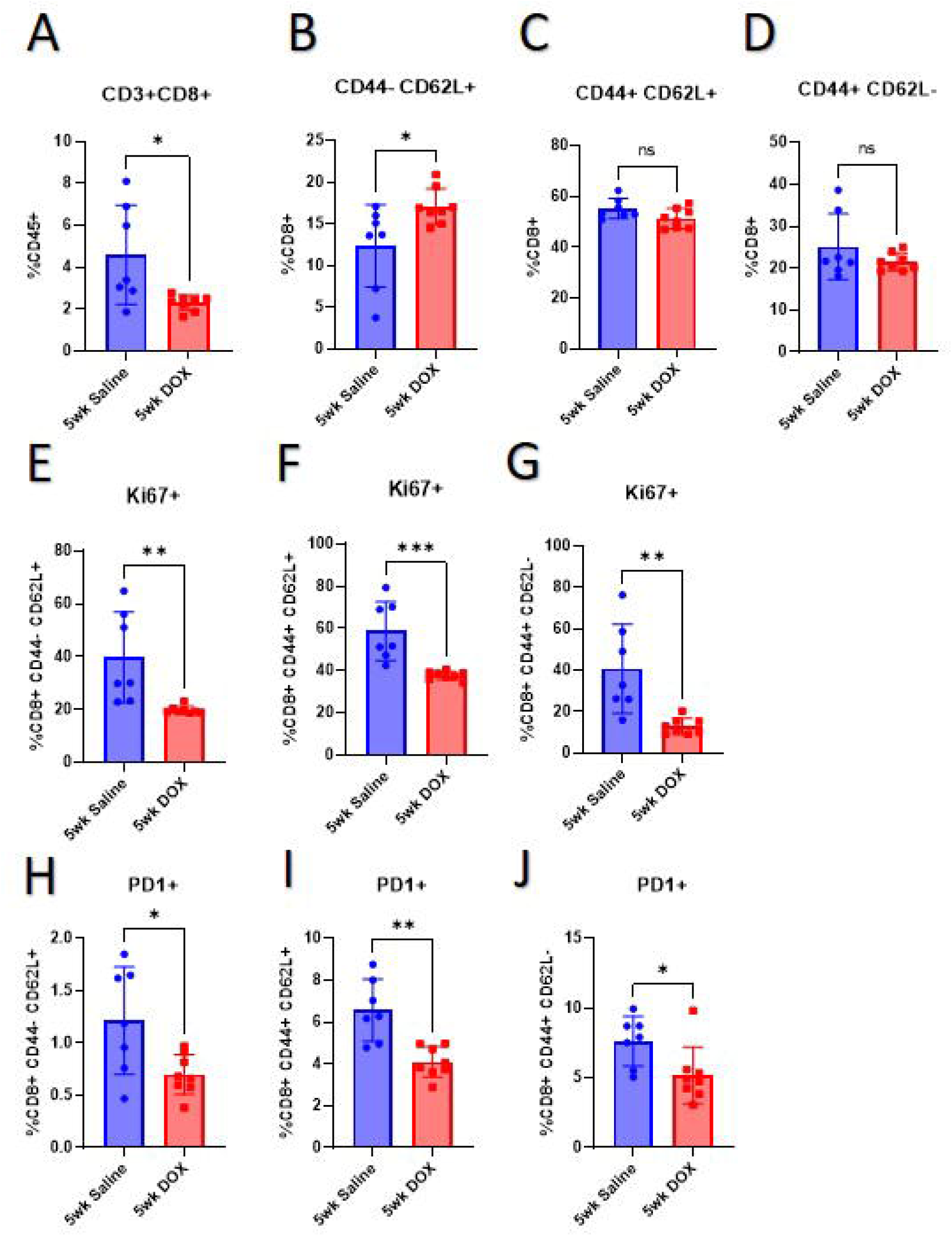
DOX induced changes in CD8^+^ T cell populations five weeks after treatment. Bar graphs representing frequencies of **A)** mature CD3^+^CD8^+^, **B)** naïve CD8^+^ T cells, **C)** central memory CD8^+^ T cells and **D)** effector memory CD8^+^ T cells; Bar graphs representing frequencies of **E)** Ki67^+^ in CD3^+^CD8^+^naïve T cells, **F)** Ki67^+^ in central memory CD8^+^ T cells and **G)** Ki67^+^ in effector memory CD8^+^ T cells; Bar graphs representing frequencies of **H)** PD1^+^ in naïve CD8^+^ T cells, **I)** PD1^+^ in central memory CD8^+^T cells and **J)** PD1^+^ in effector memory CD8^+^ T cells. n = 7-8 mice/group. ns *p* > 0.05, \**p* < 0.05, \*\**p* < 0.01, and \*\*\**p* < 0.001 vs. the one-week saline control group (analyzed by unpaired t-test).

To address how T cell subsets could be altered at this timepoint, we examined thymus gene expression for markers of thymic development, cytokines and markers associated with cellular senescence. The gene expression of markers of thymus development, of cytokines or of most markers (Fig 8 and Sup. Fig. 7) of senescence (Fig 8) did not reveal any significant alterations but there was a persistent elevation in *p21^Cip1^* gene expression (Fig. 8G and Sup. Fig. 3). No significant changes were observed in markers of thymic development including *Foxn1* (Fig. 8A) and *Pax1* (Fig. 8B). Expression of the cytokines *Il6* (Fig. 8C), *Ifny* (Fig. 8D), *Il7* (Fig. 8E), *Il17* (Fig. 8F), *Il2* (Sup. Fig 7A), *Tnf*α (Sup. Fig. 7B), and *Il4* (Sup. Fig. 7C) also remained unaltered by DOX treatment. Markers of senescence including *p16, p53,* and *p19* (Sup. Fig. 8) were not significantly altered, but there was a persistent elevation of *p21^Cip1^* gene expression (Fig. 8G).

**Figure 8:**
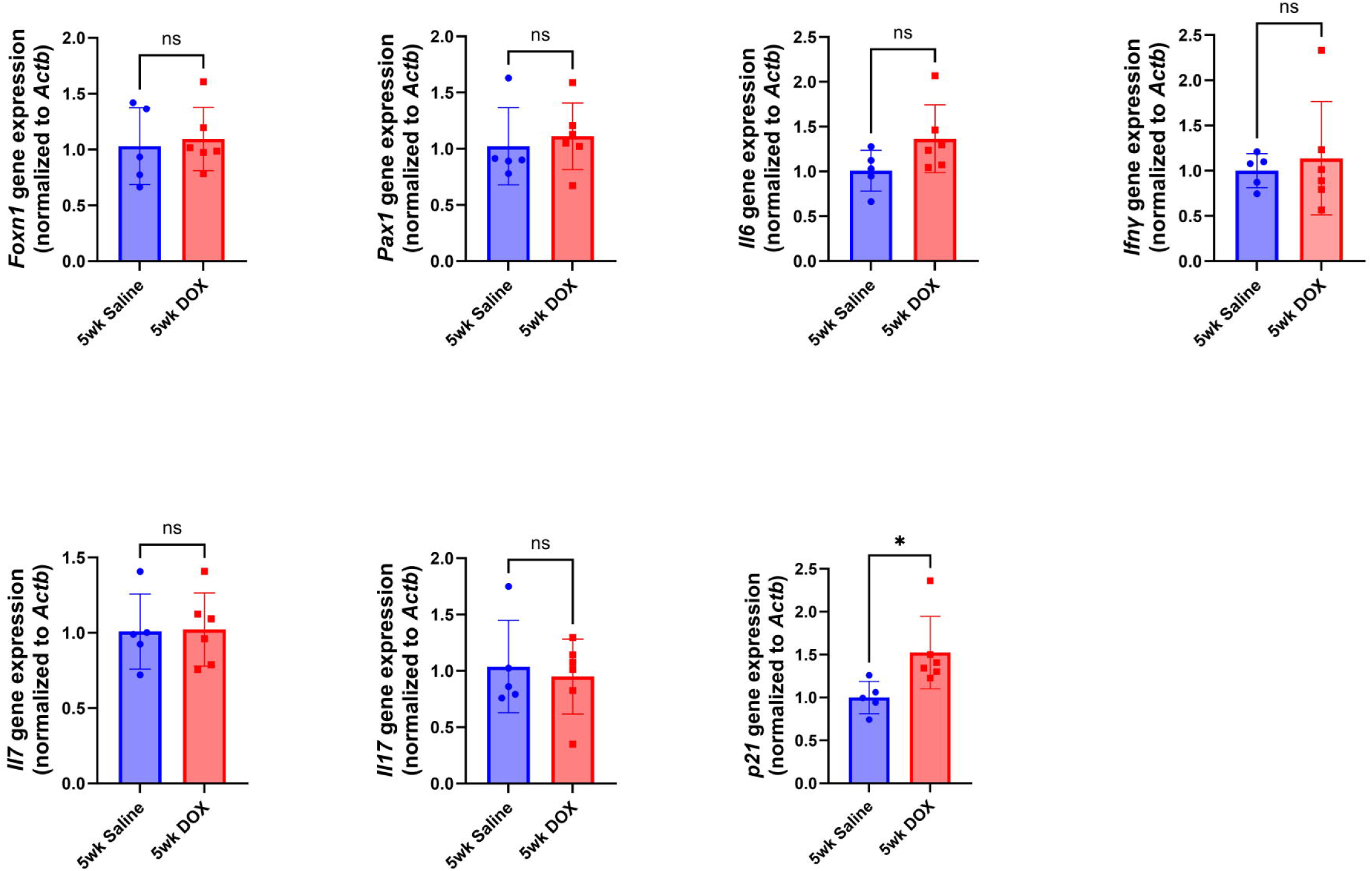
Delayed effects of DOX on the cytokines and senescence marker in thymus: Thymus were harvested from male mice five weeks following the administration of 4 mg/kg/week DOX or an equivalent volume of sterile saline for three weeks (n = 5–8 per group). Following the extraction of total RNA, the mRNA expression of **A)** *Foxon1* and **B)** *Pax1,* **C)** *Il6*, **D)** *Ifn*γ **E)** *Il7*, **F)** *Il17*, and **G), H)** *p21^Cip1^* at five weeks, was determined by qRT-PCR. Values were normalized to Actb expressed relative to saline-treated male mice. Values are shown as the means ± SEMs. The statistical significance of pairwise comparisons was determined by unpaired t-test).

### Histopathology Analysis

To examine possible pathological lesions 5 weeks after DOX administration, a total of 12 samples (6 control and 6 DOX-exposed animals) were evaluated, all of which demonstrated rare, minimal microscopic pathology. All 12 animals demonstrate rare individual cell necrosis, one animal (8%) demonstrated rare small foci of hemorrhage, and one (8%) demonstrated rare foci of cystic degeneration (consistent with physiologic thymic involution). No inflammation or edema was observed in any evaluated tissue (Fig. 9A-F). Thymocytes appeared to be similarly sized between the two groups, suggesting that the increase in thymus weight is hyperplastic, not hypertrophic (Fig. 9C and 9F). The observed findings of rare individual cell necrosis, hemorrhage, and cystic degeneration are interpreted as normal background lesions and are not considered histologically significant.

**Figure 9.**
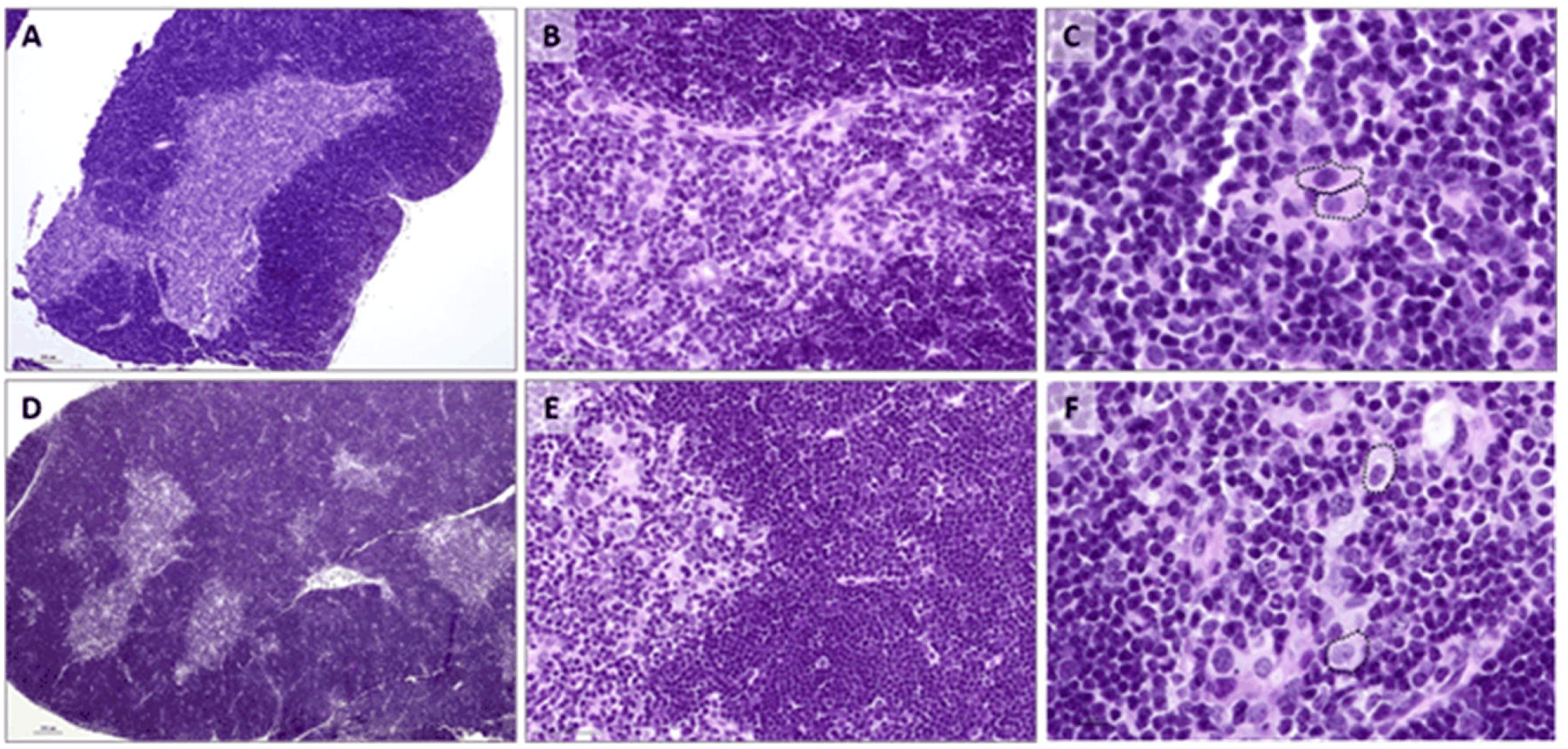
Representative thymic histology, H&E-stained sections. A-C = Saline treated; D-F = DOX treated. Across all animals, no significant histologic findings or differences were identified between the two study groups. Rare and minimal foci of hemorrhage and rare individually necrotic cells were seen. Note the similarly sized thymocytes (C&F, black dashed outline). A and D = 40X; scale bar = 100 µm, B and E = 200X; scale bar = 50 µm, C and F = 400X; scale bar = 10 µm.

## Discussion

Young adult cancer survivors experience immune dysregulation as a result of their cancer treatment. Cancer treatments such as chemotherapy and radiation therapy can weaken the immune system, making survivors more susceptible to infections and autoimmune diseases [55]. Chemotherapy not only targets cancer cells, but also affects rapidly dividing cells throughout the body, including immune cells. This can lead to a temporary suppression or dysregulation of the immune system, leaving survivors vulnerable to infections during treatment and potentially for years after treatment has ended [56]. T cells, a type of immune cell, play a crucial role in the immune system’s response to cancer and infections. However, cancer treatments can suppress T cell production or function, leaving survivors more vulnerable to infections and other immune-related complications [57]. Routine blood count tests, including measurements of immune cell counts, can provide valuable information about the immune health of cancer survivors. A decrease in immune cell count, known as leukopenia, can indicate a weakened immune system and increased susceptibility to infections. Monitoring T cell levels within the immune cell population can offer further insights into the specific components of the immune system affected by cancer and its treatments [58]. DOX administration in male juvenile mice leads to significant thymic atrophy, characterized by reductions in thymus weight, cell count, and T cell populations. This aligns with studies demonstrating the cytotoxic effects of DOX on rapidly dividing thymocytes, resulting in apoptosis and thymic involution [8, 47]. DOX induces thymic involution and decreases thymocyte numbers through the generation of reactive oxygen species and inhibition of DNA synthesis, culminating in apoptosis. Histopathological analyses confirm decreased cortical cell numbers and increased apoptosis [45]. Additionally, DOX enhances CD4^+^ T-cell responses by upregulating CD40 ligand and 4-1BB expression, which are crucial for T-cell activation. It also reduces myeloid-derived suppressor cells, thereby restoring T lymphocyte activity [8]. Chronic endurance exercise before DOX exposure can reduce oxidative stress in the thymus. However, it does not prevent thymic involution and thymocyte loss, indicating that exercise mitigates some oxidative damage but does not counteract the cytotoxic effects of DOX [45]. While the immediate effects of DOX, such as thymic atrophy and immune response alterations, are well-documented, the long-term consequences on thymic function and immune competence remain underexplored [59]. Understanding these long-term effects is crucial for developing strategies to mitigate the adverse impact of DOX on the immune system. Additionally, the decrease in thymus weight may also be influenced by alterations in the thymic microenvironment, such as changes in cytokine production and thymic epithelial cell function, induced by DOX exposure [45]. Interestingly, the delayed effects of DOX on the thymus showed a contrasting pattern, with an increase in thymus weight observed after a five-week period without drug administration. This phenomenon may reflect a compensatory response to thymic atrophy induced by DOX, characterized by increased thymic cell proliferation and expansion. However, despite the increase in thymus weight, there was a diminished occurrence of hematopoietic cells, suggesting potential alterations in thymic microenvironment and function induced by DOX exposure.

Upon examining the immediate impact of DOX on CD4^+^ T cells, we observed no significant changes in the frequency of mature CD3^+^CD4^+^ T cells, but there were elevated frequencies of exhaustion markers, specifically PD1 expression, in naïve CD4^+^ T cells within the DOX-treated group. Interestingly, central memory CD4^+^ T cells showed a significant decrease in frequency, while effector memory CD4^+^ T cells exhibited lower proliferation and exhaustion frequencies in the DOX group compared to saline. The elevation of exhaustion markers in naïve CD4^+^ T cells and the decrease in central memory T cells suggest a disruption in T cell maturation processes. The lower proliferation and exhaustion frequencies in effector memory CD4^+^ T cells may reflect a compensatory response to DOX-induced stress. Similarly, the immediate effects of DOX on CD8^+^ T cells revealed a significant decrease in the frequency of mature CD3^+^CD8^+^ T cells and an increase in the frequency of naïve CD8^+^ T cells. Notably, there was an elevation in exhaustion markers in naïve CD8^+^ T cells within the DOX group. Despite no significant changes in the frequency of central memory and effector memory CD8^+^ T cells, there was a reduction in proliferation and exhaustion frequencies in these subsets following DOX treatment. The reduction in proliferation and exhaustion frequencies in central memory and effector memory CD8^+^ T cells could signify a protective mechanism against DOX-induced damage.

After the conclusion of chemotherapy, thymic hyperplasia becomes evident, showing no apparent correlation with the extent of lymphocyte depletion. Instead, it emerges as a prevalent phenomenon following the cessation of chemotherapy [4, 15]. Numerous case reports have detailed occurrences of thymic hyperplasia following adjuvant chemotherapy in individuals diagnosed with breast cancer [4, 15]. Considering the observed clinical thymic hyperplasia, our focus turned to understanding the delayed impact of DOX on the thymus in adult mice that were pre-exposed to DOX as juveniles, along with an exploration of changes within the immune cell population and subpopulations. Despite no apparent correlation with the extent of lymphocyte depletion, an increase in the thymus size was evident in DOX-treated male mice, suggesting a complex interplay between thymic responses and chemotherapy. This phenomenon aligns with numerous case reports detailing occurrences of thymic hyperplasia following chemotherapy in cancer survivors [19–21, 62, 63], but it has never been reported in a preclinical animal model. Interestingly, while there was no significant difference in the frequency of immune cells (CD45^+^), there was a diminished occurrence of hematopoietic cells in the delayed phase, suggesting a potential long-term effect on thymic cellularity. The delayed impact of DOX on immune cell subsets revealed significant alterations in the proliferation and exhaustion patterns of CD4^+^ T cells. While mature CD3^+^CD4^+^ cells and central memory CD4^+^ T cells exhibited significant decreases in frequency, no significant changes were observed in naïve CD4^+^ T cells. DOX treatment led to a significant decrease in the proliferation and exhaustion of naïve CD4^+^ T cells, suggesting immunosuppressive effects [64, 65]. Similar effects were observed in CD8^+^ T cell subsets, with a significant decrease in mature CD3^+^CD8^+^ T cells and a notable increase in naïve CD8^+^ T cells following DOX treatment. While the frequencies of central memory and effector CD8^+^ T cells showed no significant changes, decreases in proliferation and exhaustion frequencies were observed across all CD8^+^ T cell subsets in the DOX group compared to saline. While central memory and effector CD8^+^ T cell frequencies remained unchanged, decreases in proliferation and exhaustion frequencies were evident across all CD8^+^ T cell subsets in the DOX group compared to the saline group [65, 66].The treatment with DOX significantly decreased both the proliferation (indicated by reduced Ki67 expression) and the initial activation markers of naïve CD4^+^ T cells, notably PD1 expression. This suggests that DOX may suppress immune responses by inhibiting the activation and proliferation of these cells, reducing their progression towards exhaustion. Consequently, DOX-induced selective targeting of specific CD4^+^ T cell populations may contribute to broader immunosuppressive effects.

The roles of Foxn1 and Pax1 in thymic development are essential, with Foxn1 facilitating the transcription of genes critical for thymus organogenesis and TEC differentiation, and Pax1 creating a microenvironment for T cell maturation [49]. In DOX-treated male mice, increased expression of these transcription factors suggests adaptive mechanisms aimed at maintaining thymic integrity and function under stress. This indicates that despite the chemotherapeutic challenge, the thymus attempts to preserve its epithelial microenvironment and support T cell development. The cytokine network within the thymus also adapts to DOX treatment, evidenced by decreased *Il6* expression, which is vital for thymocyte proliferation and differentiation [48]. This reduction could impair thymic function, leading to degeneration and compromised T cell development [67, 68]. Conversely, increased *Ifn-*γ suggests a shift towards a Th1-biased immune response [70]. Additionally, the upregulation of *Il7* indicates a compensatory effort to support thymocyte survival and proliferation despite thymic degeneration [67, 70]. However, the decrease in *Il17* expression could further contribute to thymic atrophy and impaired T cell development, complicating the immune regulatory landscape under DOX treatment [67, 71]. These findings highlight the delicate balance of cytokine signaling and transcriptional regulation necessary for maintaining thymic function under chemotherapeutic stress.

DOX induces DNA double-strand breaks, which activate the ATM/ATR/Chk1/Chk2 pathway, leading to the stabilization and activation of p53. Activated p53 then upregulates p21 transcription, causing cell cycle arrest and promoting senescence [73–76]. In the thymus, DOX-induced DNA damage triggers a p53-dependent response, leading to increased expression of p21 and other cell cycle arrest proteins [74, 76]. DOX treatment leads to increased *p21^Cip1^* expression in the thymus, which is associated with thymic involution and impaired T cell development. This can result in a reduced T cell repertoire and compromised immune function, increasing the risk of infections and diseases in aging individuals [8, 76]. The persistent high expression of *p21^Cip1^*following DOX treatment suggests a sustained senescence response, which can disrupt thymic architecture and function, further impacting T cell maturation [8, 76]. The findings presented in this study have important implications for understanding the impact of DOX on thymus function and immune responses, particularly in childhood cancer survivors. The observed alterations in thymic morphology and T cell populations may contribute to immunosenescence, potentially affecting the long-term immune health of survivors. Further research is warranted to elucidate the mechanisms underlying DOX-associated thymus changes and their effects on immune function, with the ultimate goal of optimizing treatment strategies and improving outcomes for childhood cancer survivors.

## Author Contributions

B.G.; Conceptualization, Data curation, Formal analysis, Methodology, Writing – original draft

K.J.V.D; Conceptualization, Data curation, Formal analysis, Methodology, Writing – original draft

M.K.O.G; Data curation, Formal analysis, Methodology, Project Administration

D.M.S; Data curation, Formal analysis, Methodology

M.R.D; Data curation, Formal analysis, Methodology

D.S; Data curation, Formal analysis, Methodology

K.T.S; Conceptualization, Writing – reviewing and editing

C.D.C; Conceptualization, Funding acquisition, Supervision, Writing – reviewing and editing

B.N.Z; Conceptualization, Funding acquisition, Supervision, Writing – reviewing and editing

