## Supplementary Figures for "Divergent Immediate and Delayed Effects of Juvenile Exposure to Doxorubicin on the Thymus in C57BL/6 Mice"

**Figure 1: A)** immediate effects of DOX on the body weight; **B)** recorded longitudinally body weight for five week following the third dose of DOX; **C)** immediate effects of DOX on the thymus weight (normalized to tibial length) and **D)** immediate effects of DOX on the thymus weight (normalized to body weight)

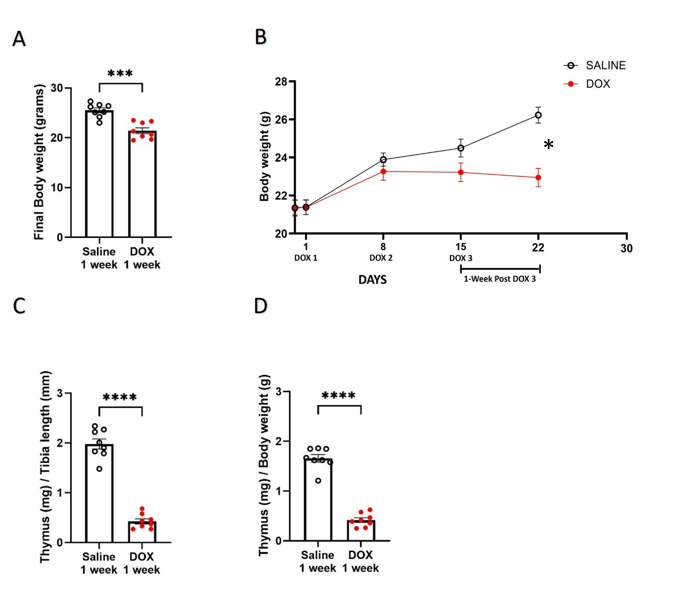

**Figure 2:** Thymocyte gating strategy.

Representative flow cytometry pseduocolor plots for flow cytometry data shown in Figure 1.

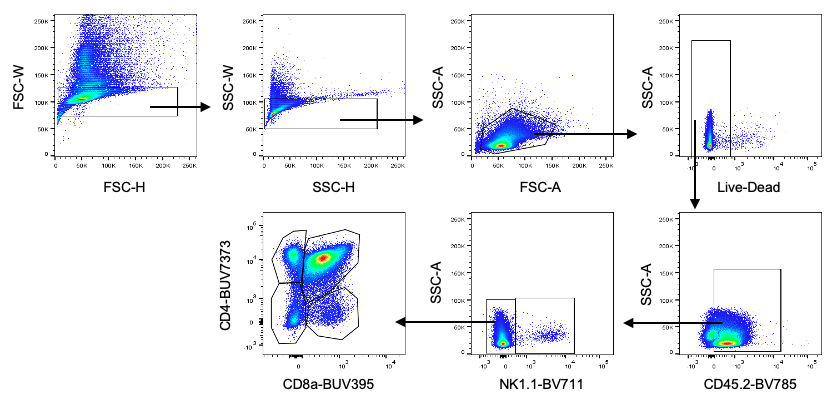

**Figure 3:** Mature T cell gating Strategy.

Representative flow cytometry pseduocolor plots for flow cytometry data shown in Figure 2 and 3.

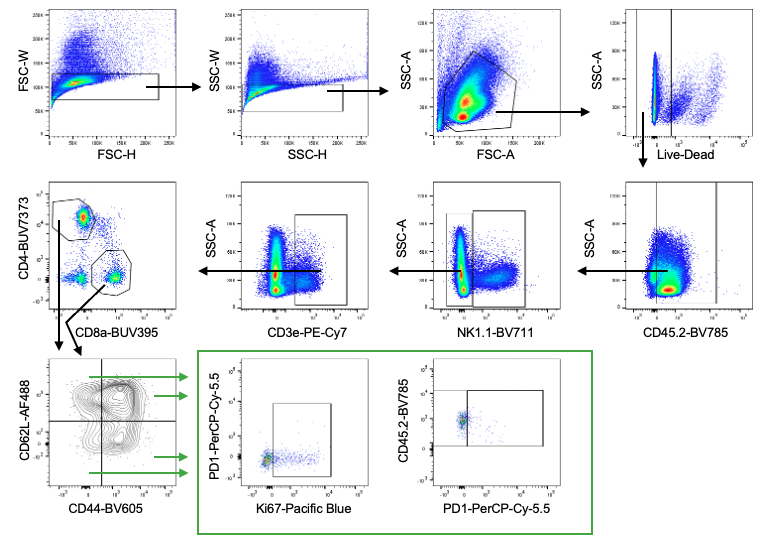

**Figure 4:** Thymus were harvested from male mice one weeks following the administration of 4 mg/kg/week DOX or an equivalent volume of sterile saline for three weeks (n=5–8 per group). Following the extraction of total RNA, the mRNA expression of **A)** *Il2*, **B)** *Tnfα* **C)** *Il4*, was determined by real-time PCR. Values were normalized to *Actb* and expressed relative to saline-treated male mice. Values are shown as the means ± SEMs. The statistical significance of pairwise comparisons was determined by unpaired t-test).

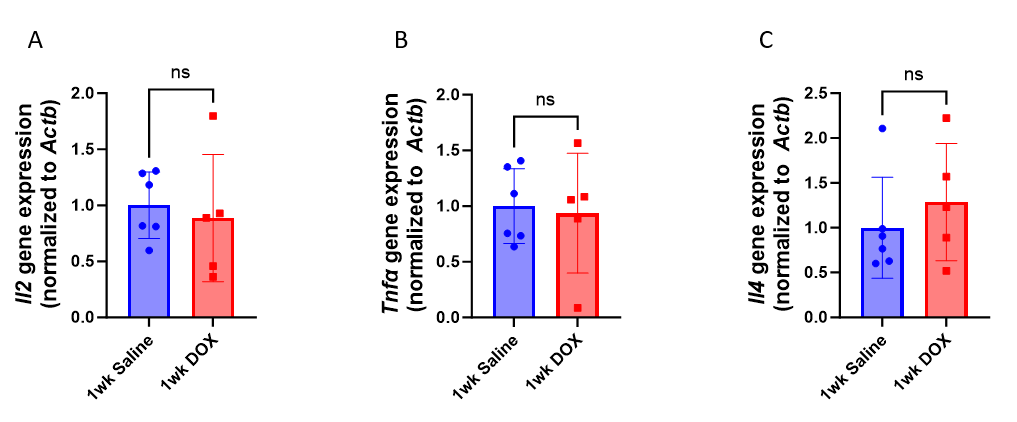

**Figure 5:**  **Effects of DOX on the senescence marker in thymus:**

Thymus were harvested from male mice one week following the administration of 4 mg/kg/week DOX or an equivalent volume of sterile saline (n=5–8 per group). Following the extraction of total RNA, the mRNA expression of **A)** *p16*, **B)** *p53,* **C)** *p19* senescence marker at one week was determined by real-time PCR. Values were normalized to *Actb* and expressed relative to saline-treated male mice. Values are shown as the means ± SEMs. The statistical significance of pairwise comparisons was determined by unpaired t-test.

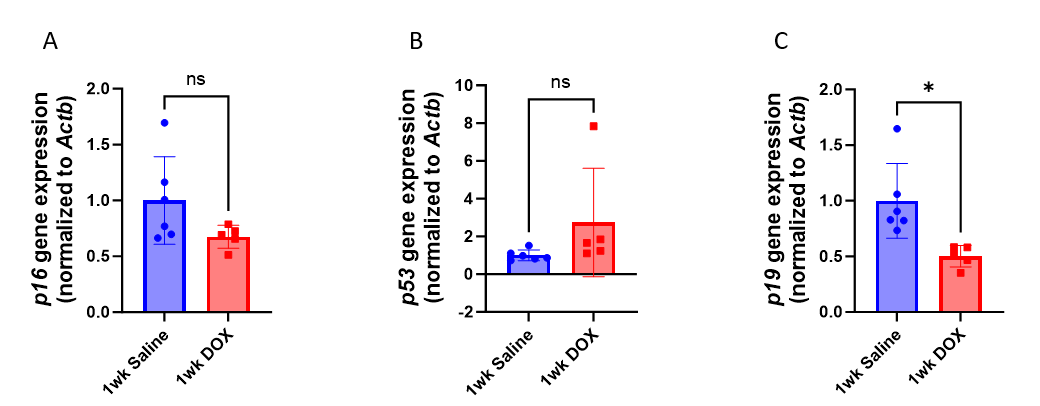

**Figure 6: A)** delayed effects of DOX on the body weight; **B)** recorded longitudinally body weight for five week following the third dose of DOX; **C)** delayed effects of DOX on the thymus weight (normalized to tibial length); and **D)** delayed effects of DOX on the thymus weight (normalized to body weight).

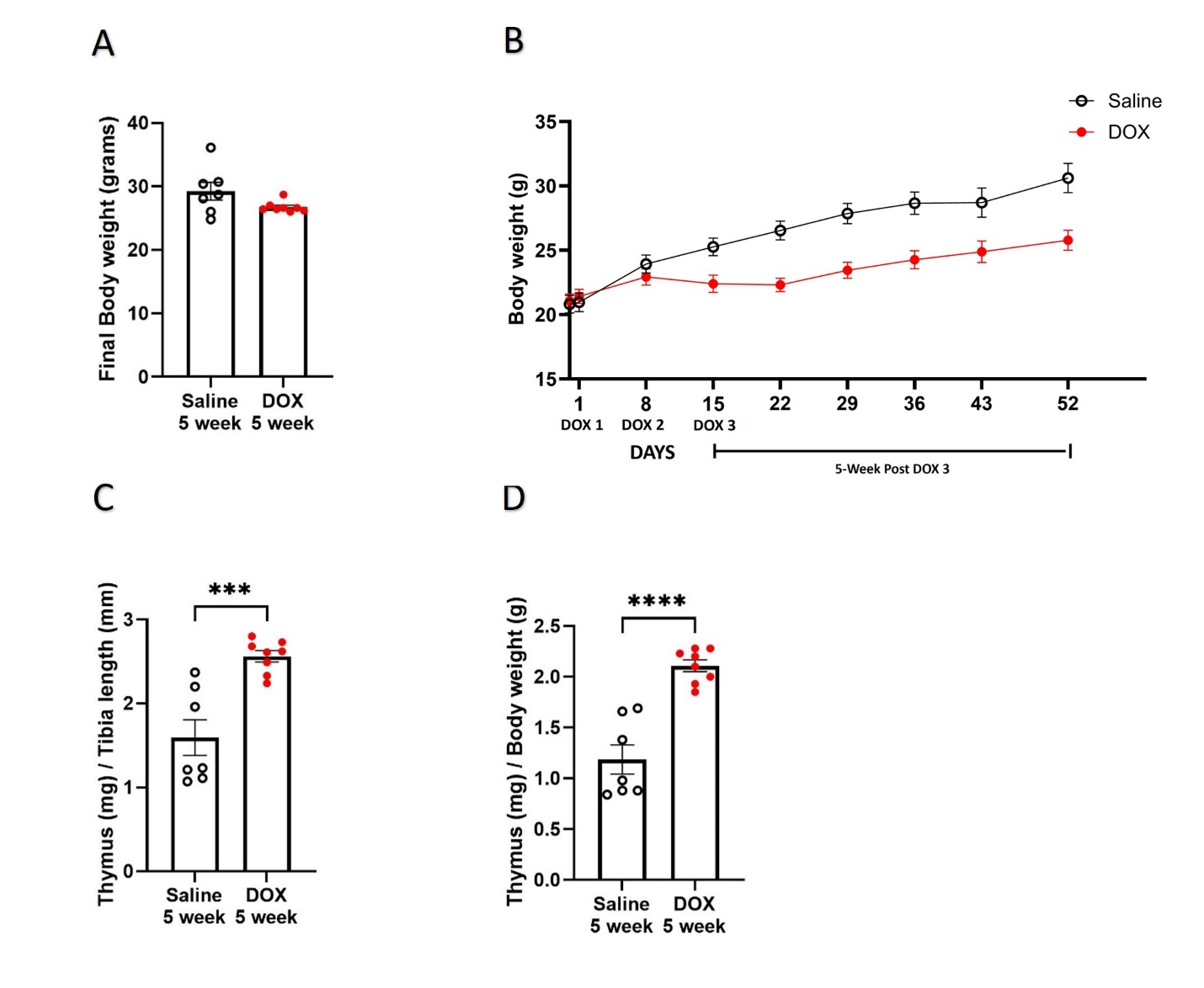

*

**Figure 7:** Thymus were harvested from male mice five weeks following the administration of 4 mg/kg/week DOX or an equivalent volume of sterile saline (n=5–8 per group). Following the extraction of total RNA, the mRNA expression of **A)** *Il2*, **B)** *Tnfα* **C)** *Il4*, was determined by real-time PCR. Values were normalized to *Actb* and expressed relative to saline-treated male mice. Values are shown as the means ± SEMs. The statistical significance of pairwise comparisons was determined by unpaired t-test.

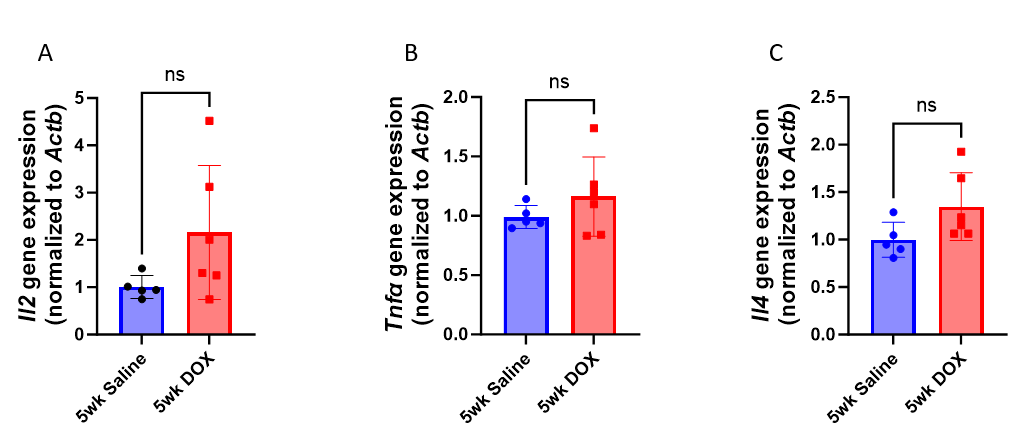

**Figure 8:**  **Effects of DOX on the senescence marker in thymus:**

Thymus were harvested from male mice five week following the administration of 4 mg/kg/week DOX or an equivalent volume of sterile saline for three weeks (n=5–8 per group). Following the extraction of total RNA, the mRNA expression of **A)** *p16*, **B)** *p53,* **C)** *p19* senescence marker at five week was determined by real-time PCR. Values were normalized to *Actb* and expressed relative to saline-treated male mice. Values are shown as the means ± SEMs. The statistical significance of pairwise comparisons was determined by unpaired t-test.

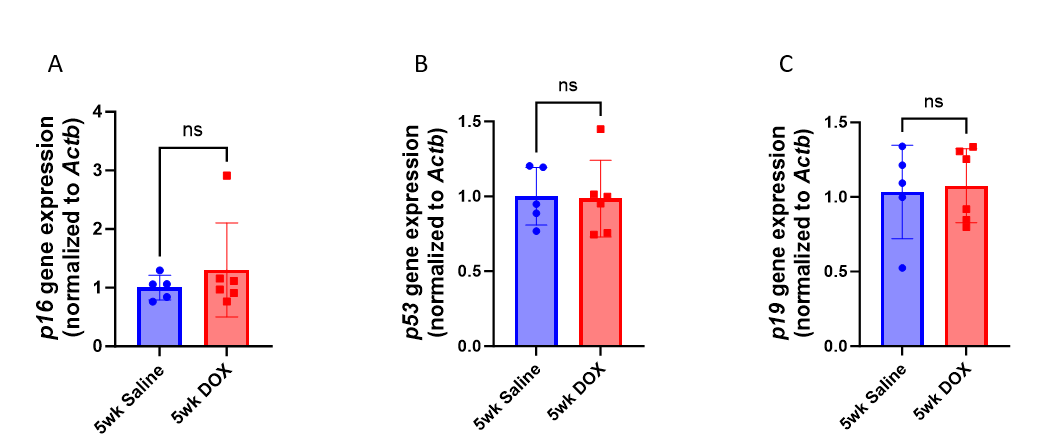
